# Enzymatic synthesis of fatty acid amides using microbial lipids as acyl group-donors and their biological activities

**DOI:** 10.1101/2020.09.13.295113

**Authors:** Hatim A. El-Baz, Ahmed M. Elazzazy, Tamer S. Saleh, Panagiotis Dritsas, Jazem A. Mahyoub, Mohammed N. Baeshen, Hekmat R. Madian, Mohammed Alkhaled, George Aggelis

## Abstract

Fatty acid amides (FAAs) are of great interest due to their broad industrial applications. They can be synthesized enzymatically with many advantages over chemical synthesis. In this study, the fatty acid moieties of lipids of *Cunninghamella echinulata* ATHUM 4411, *Umbelopsis isabellina* ATHUM 2935, *Nannochloropsis gaditana* CCAP 849/5, Olive oil and an eicosapentaenoic acid (EPA) concentrate were converted into their fatty acid methyl esters and used in the FAA (i.e. ethylene diamine amides) enzymatic synthesis, using lipases as biocatalysts. The FAA synthesis, monitored using *in situ* NMR, FT-IR and thin-layer chromatography, was catalyzed efficiently by the immobilized *Candida rugosa* lipase. The synthesized FAAs exhibited a significant antimicrobial activity, especially those containing oleic acid in high proportions (i.e. derived from Olive oil and *U. isabellina* oil), against several human pathogenic microorganisms, insecticidal activity against yellow fever mosquito, especially those of *C. echinulata* containing gamma linolenic acid, and anti-cancer properties against SKOV-3 ovarian cancer cell line, especially those containing EPA in their structures (i.e. EPA concentrate and *N. gaditana* oil). We conclude that FAAs can be efficiently synthesized using microbial oils of different fatty acid composition and used in specific biological applications.

## Introduction

Fatty acid amides (FAAs) are organic compounds formed from a fatty acid (FA) and an amine, such as ethanolamine or an amino acid. FAAs can be synthesized from alkanolamine and a fatty acyl donor, such as a free FA or a FA alkyl ester, by chemical or enzymatic esterification or transesterification methods [1, 2]. The enzymatic synthesis of FAAs can be performed using lipases [3, 4], or aminoacylases [5]. FAAs are of considerable interest due to their wide-ranging industrial applications in the production of lubricants, detergents, shampoo, cosmetics and surfactant formulations [6, 7]. In addition, FAAs, demonstrating a potent antimicrobial activity against Gram-positive and Gram-negative bacteria [8] and possessing beneficial anti-inflammatory properties [9], provide an exciting opportunity to produce new medicines and nutraceuticals with applications in the treatment of several human diseases and in human nutrition [7, 10].

There are different sources of FAs, such as common plant oils and animal fats, which can be used in amide synthesis. Alternatively, microbial lipids, so called Single Cell Oils (SCOs), derived from microalgae and fungi, which do not compete with the food supply chain, could be considered for this purpose. Microalgae and fungi are on the forefront of biotechnological interest due to their ability to produce SCOs rich in polyunsaturated fatty acids (PUFAs) of medical and nutritional interest [11–17]. The high PUFA content of the aforementioned lipids offers an additional interest in their use as acyl group-donors in FAA synthesis, since several reports demonstrate that PUFAs or compounds containing PUFA moieties in their molecule exhibited interesting biological activities [18–20]. Among microalgae, *Nannochloropsis* is a prominent genus that include species able to efficiently grow under non-aseptic conditions and accumulate lipids rich in PUFAs, such as eicosapentaenoic acid (EPA) [21, 22]. As for fungi, genera belonging to Mucoromycota (including *Mucor, Rhizopus, Umbelopsis, Lichtheimia, Cunninghamella* and *Mortierella*) are well known for their ability to synthesize PUFAs, especially gamma linolenic acid (GLA), which is of great pharmaceutical interest due to its anticancer properties, while it has been used to improve premenstrual tension and various skin diseases [13, 17]. Especially, *Cunninghamella echinulata* is an important GLA producer [13, 19, 23], while *Umbelopsis isabellina* is regarded as a promising SCO producer, being able to accumulate lipids in high percentages, though less rich in GLA [16, 24–26].

The aim of this study was to produce through enzymatic synthesis FAAs using as acyl group-donors SCOs of different FA composition, such as those derived from the fungi *Umbelopsis isabellina* (containing oleic acid in high percentage and GLA in low percentage) and *Cunninghamella echinulata* (containing GLA in high percentages) and the microalga *Nannochloropsis gaditana* (containing EPA in high percentages). The biological activity of the above FAAs was tested against important human pathogens, the larvae of *Aedes aegypti* and the SKOV-3 cancer cell line and compared with that of FAAs produced using as acyl group-donors Olive oil (containing oleic acid in very high percentages) and an EPA concentrate (i.e. a fish oil derivative containing EPA in very high percentages). We concluded that FAAs can be efficiently produced using lipids of microbial origin and employed as bioactive compounds in various biological applications depending on their FA composition.

## Materials and methods

### Biological materials and culture conditions

The fungal strains *Cunninghamella echinulata* ATHUM 4411 and *Umbelopsis isabellina* ATHUM 2935 (culture collection of National and Kapodistrian University of Athens, Greece) were maintained on potato dextrose agar (PDA) (Biolab Zrt, Budapest, Hungary) at 7 ± 1 °C. The microalga *Nannochloropsis gaditana* CCAP 849/5 was maintained in 250-mL conical flasks containing 50 mL of artificial sea water (ASW) at 25 ± 1 °C. All cultures were regularly sub-cultured.

Cultures of *C. echinulata* and *U. isabellina* were performed in 250-mL Erlenmeyer flasks containing 50 mL of a culture medium with the following composition (in g/L): glucose (AppliChem, Darmstadt, Germany), 60.0; KH_2_PO_4_ (AppliChem), 12.0; Na_2_HPO_4_ (AppliChem), 12.0; CaCl2 2H_2_O (Carlo Erba, Rodano, Italy), 0.1; CuSO_4_ 5H_2_ O (BDH, Poole, England), 0.0001; Co(NO_3_) 6H_2_O (Merck, Darmstadt, Germany), 0.0001; MnSO_4_ 5H_2_O (Fluka, Steinheim, Germany), 0.0001; ZnSO_4_ 7H_2_O (Merck), 0.001 and FeCl_3_ 6H_2_O (BDH), 0.08. The medium was limited in nitrogen with yeast extract (Conda, Madrid, Spain) at 3.0 g/L being the sole nitrogen source. Yeast extract was also served as source of magnesium and ferrum according to Bellou et al. [13]. The flasks were sterilized at 121 °C for 20 min and inoculated with 1 mL of spore suspension containing 10^7^ fungal spores produced on PDA cultures for 5 days at 28 °C. Incubation took place in an orbital shaker (ZHICHENG ZHWY 211C, Shanghai, China) at temperature 28 ± 1 °C and an agitation rate of 180 rpm. pH after sterilization was 6.5 ± 0.5 and remained practically stable during cultivation.

A modified ASW described in Dourou et al. [22] was used as growth medium for *N. gaditana*. Prior to sterilization pH of ASW was calibrated at 8.5 ± 0.5 through the addition of 2 M NaOH (Merck) solution. Microalgal cultures were performed in a laboratory-made glass bioreactor of total volume 8.7 L and working volume 5.0 L, served as an open-pond simulating reactor (OPSR) [22]. Initially, the reactor was washed with 70% ethanol and filled with 4.5 L sterilized (at 121 °C for 20 min) ASW medium. The reactor was inoculated with 500 mL of a fresh inoculum containing 10^5^ cells/mL, and incubated at temperature 25 ± 1 °C under constant illumination of 300 μmol m^−2^ s^−1^ supplied by linear fluorescent day light tubes T5, 8W, 65OOk, G5. OPSR cultures were performed at temperature 25 ± 1 °C and pH 8.5 ± 0.5, which was automatically controlled. Agitation was achieved through the use of a circulator, in the entryway of which natural air was provided to the culture with a gas flow rate of 30 L/h. Illumination of 245 μmol m^−2^ s^−1^ was provided by 8 W fluorescent lamps, which were placed at a distance of 20 cm above the culture surface.

### Cell mass harvesting

Fungal mycelia were harvested by filtration through Whatman No. 1 paper. Microalgal cell mass was harvested by centrifugation at 24,000 g for 15 min at 4 °C (Heraeus, Biofuge Stratus, Osterode, Germany). In both cases, the collected biomass was washed twice with distilled water, dried at 80 °C until constant weight and gravimetrically determined.

### Lipid extraction and purification

Microbial lipids were extracted in chloroform: methanol (2:1, v/v) (Sigma-Aldrich) following the Folch et al. [27] method. The extracts were filtrated through Whatman No. 1 paper and washed with a KCl (Sigma-Aldrich) 0.88 % (w/v) solution to remove non-lipid components. Subsequently, the solvents were dried over anhydrous Na_2_SO_4_ (Sigma-Aldrich) and evaporated under vacuum using a Rotavapor R-20 device (BUCHI, Flawil, Switzerland). The total cellular lipids (L) were gravimetrically determined and expressed as a percentage on dry cell mass (L/x, %).

### Fatty acid methyl esters preparation and gas chromatography analysis

The FA moieties of lipids (i.e. approx. 100 mg of microbial oils or Olive oil or EPA concentrate produced as above described) were converted into their fatty acid methyl esters (FAMEs) in a two-stage reaction in accordance with the AFNOR [28] method in order to avoid trans-isomerization. Briefly, in the first stage the FAs that are esterified with glycerol were converted into FAMEs and the free FAs (if present) were converted into sodium soaps in a sodium methoxide solution under reflux. Following, the resulting soaps were also converted into FAMEs after adding an acetyl chloride solution in excess in the above mixture. The reaction was stopped by adding water and the FAMEs were extracted in 6 mL hexane (Fluka). Finally, the organic phase was removed under vacuum and the FAME preparation was stored in dark under an argon atmosphere.

FAME mixtures were analysed in a gas chromatography device (Agilent 7890A device, Agilent Technologies, Shanghai, China), equipped with a flame ionization detector (working at 280 °C) and a HP-88 (J&W Scientific) column (60 m × 0.32 mm). Carrier gas was helium at a flow rate 1 mL/min and the analysis was run at 200 °C. Peaks of FAMEs were identified through comparison to authentic standards.

### Free fatty acid preparation

For glycerides cleavage, 1 g of lipids was saponified in 10 mL KOH 1N ethanol solution (95%) under reflux for 1 h and 45 min. The mixture was acidified with 10 mL HCl 4 N solution and the free FAs were extracted three-times with 5 mL hexane. The organic phase was washed with distilled water until the washes were neutral and dried over anhydrous Na_2_SO_4_ (Sigma). Finally, the organic phase was removed under vacuum and the FFA preparation was stored under an Argon atmosphere.

### Enzymatic synthesis of amides

The lipase-catalyzed synthesis of amides was carried out in 50-mL Erlenmeyer flasks in nearly anhydrous media using 100 mg Novozym 435 lipase (i.e. immobilized *C. antarctica* lipase, enzymatic activity □2 Units/mg) or 100 mg lipase from *C. rugosa* (immobilized, enzymatic activity □2 Units/mg), both purchased from Sigma Aldrich Co., St. Louis, MO, as biocatalysts. The reaction was carried out in an orbital shaker at 40 ± 1 °C, 90 rpm, in 25 mL acetone (Sigma Aldrich Co.), with ethylene diamine (Acros Organics, Thermo Fisher Scientific, Waltham, MA) and FAMEs or FFAs (produced as above) as substrates, used at different molar ratios. After several preliminary experiments FAME preparations were selected as substrate. The reaction lasted until the FAME substrate was exhausted, as evidenced by TLC (see below), usually after 18 h of incubation. The reaction mixture was then filtered, the solvent was removed from the filtrate by evaporation under reduced pressure and the reaction residue is partitioned in dichloromethane and distilled water (20 mL each). The organic layer, containing the synthesized amide, was washed with saturated aqueous NaCl (Sigma Aldrich Co.) (10 mL), dried over MgSO_4_ (Sigma Aldrich Co.), gravity-filtered and the solvent removed under reduced pressure to get the crude product.

### Monitoring the evolution of the reaction and product characterization Thin-layer chromatography and FT-IR

The reaction was monitored by thin-layer chromatography (TLC) performed on precoated Merck 60 GF254 silica gel plates (Merck, US) with a fluorescent indicator, and visualized under ultraviolet irradiation at 254 and 360 nm. A mixture of n-hexane: ethyl acetate (1:4) was used as eluent and the progress of the reaction monitored until the disappearance of FAME spot in the reaction mixture.

FT-IR spectra for FAMEs and the FAA products were recorded on a Smart iTR, which is an ultrahigh-performance, versatile attenuated total reflectance sampling accessory on the Nicolet iS10 FT-IR spectrometer (Thermo Fisher Scientific). FT-IR spectra were used to confirm the formation of FAAs from FAMEs, by detecting the formation of the amidic carbonyl group.

### Quantitative determination of the reaction yields through *in situ* NMR monitoring

The percent conversion of FAMEs to amides was calculated during the reaction via *in situ* NMR monitoring. In detail, the protons of methyl group of FAMEs, which are present at δ 3.56 ppm, were assigned and the progress of the reaction of FAMEs with ethylene diamine was monitored by ^1^H NMR at regular intervals of 6 h. This was achieved by drawing a sample from the reaction mixture using a 1mL syringe connected with a syringe filter of 0.22 μm pore size, followed by evaporation of the solvent and dissolution of the residue in CDCl_3_. The ^1^H NMR was noted a new singlet signal at δ 3.91 ppm which matched the -CH_2_-CH_2_-of the amine used and grew concurrently with a decline in the intensity of the methyl of ester group signals. The latter signals disappeared after 24 h for ratio of FAME: amine 1:5 indicating 100% conversion. Therefore, we succeed to calculate the % conversion via integrations of the peaks originated one from the product (p) and the other from the reactant (r) according to the formula:

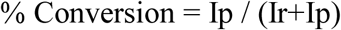

Where Ip is the integration of the signal of the product and Ir is the integration of the signal of the reactant.

The physical properties and spectral data of the prepared diamides are described below. Diamide of *Cunninghamella echinulata* methyl esters: Viscous liquid; black; IR (ν_max_, cm^−1^): 3266 (NH), 2924, 2853 (CH_2_), 1651 (CO amidic); Diamide of *Umbelopsis isabellina* methyl esters: Liquid; dark brown; IR (ν_max_, cm^−1^): 3276 (NH), 2924, 2854 (CH_2_), 1663 (CO amidic); Diamide of *Nannochloropsis gaditana* methyl esters: Viscous liquid; Yellow, IR (ν_max_, cm^−1^): 3266 (NH), 2925, 2854 (CH_2_), 1663 (CO amidic); Diamide of Olive oil methyl esters: Semi solid material; off-white color; IR (ν_max_, cm^−1^): 3407 (NH), 2922, 2852 (CH_2_), 1638 (CO amidic); Diamide of EPA concentrate methyl esters: Liquid material; orange color; IR (ν_max_, cm^−1^): 3291 (NH), 2959, 2925 (CH_2_), 1663 (CO amidic).

### Antimicrobial evaluation of FAAs

The antimicrobial activity of the synthesized FAAs was tested *in vitro* using agar well diffusion assay, MIC and MBC (see below) against human pathogens including the Gram-negative *Escherichia coli* ATCC 25922, *Klebsiella pneumoniae* ATCC 700603, *Pseudomonas aeruginosa* ATCC 15442, *Salmonella typhimurium* ATCC 14028 the Gram-positive bacteria, *Bacillus subtilis* ATCC 6633, MRSA *Staphylococcus aureus* ATCC 4330, *Staphylococcus aureus* ATCC 25923 and the unicellular fungus *Candida albicans* ATCC 10221.

### Agar well diffusion assay

Fresh bacterial cultures grown on nutrient agar for 20 h at 37 °C were suspended in a saline solution (0.85%, w/v) to a turbidity of 0.5 Mac-Farland standards. Then 100□μl (10^6^ CFU/mL) of each bacterial suspension was swabbed onto Mueller Hinton II Agar (MHA) plates. 6 mm diameter wells were punched on the MHA and inside the wells 100□μl of FAAs solution was poured. The plates were preincubated in a refrigerator (at T=4□°C) for 1□h and then incubated overnight at 37□°C, in order to allow the FAAs diffusion into the agar. The diameter of the inhibition zones were measured in mm using Clinical and Laboratory Standards Institute (CLSI) guidelines. Experiments were done in triplicate.

### Evaluation of minimum inhibitory concentration (MIC) and minimum bactericidal concentration (MBC) values for FAAs

MICs were determined according to the CLSI broth microdilution method [29]. One hundred microliters of the Mueller Hinton broth medium were distributed into the wells of the micro titer plates. A FAA solution (10□µL), serial diluted from stock solutions to achieve 200, 100, 50, 25, 12.5 and 6.25 µg/mL, was added to the well together with one hundred microliters of bacterial suspension (6 × 10^6^ CFU). The microwell plates were incubated at 37□°C for 24□h, then 5 μl of a resazurin solution (6.75 mg/mL) was added to each well and the plates incubated at 37 °C for another 4 h. Changes of color indicating cell viability were recorded. The bacterial growth was measured using a Bio-Rad Microplate Reader at 600 nm. MIC was determined as the lowest concentration of FAAs that inhibit visible growth of the tested microorganism. MBC was the lowest FAA concentration resulting in microbial death. It was determined by sub-culturing cells from wells that exhibited no color change to sterile MHA plates. All experiments were carried out in triplicate.

### Larval bioassay

Tests were performed on a field strain of *Aedes aegypti* raised from wild larvae, collected from Jeddah, Saudi Arabia, and maintained in the laboratory under controlled conditions of 27 ± 1 °C and 70 ± 5% R.H., with a 14:10 (L:D) photoperiod. The standard World Health Organization larval susceptibility test method was used. Treatments were carried out by exposing early 4^th^ instar larvae of *A. aegypti* to various concentrations of the tested compounds for 48 h, in groups of glass beakers containing 100 mL of a FAA solution in tap water. Five replicates per FAA concentration of 20 larvae each, and so for control trials, were set up. The larvae were given the usual larval food during these experiments. Larval mortalities were recorded at 48 h post-treatment. Log concentration-probability regression lines were drawn for the tested compounds. Statistical parameters were calculated using the method of Finney [30].

### Quantitative analysis of cell apoptosis by flow cytometry

The apoptotic activity of the SKOV-3 ovarian cancer cell line in response to tested compounds was determined by Annexin FITC, as per the manufacturer’s instructions (BD Biosciences, USA). Briefly, the SKOV-3 cells were grown in a 25-mL flask at a density of 3 × 10^5^ cells/well. The induction of apoptosis was investigated in untreated and treated SKOV-3 cells with curcuminoids at a concentration of 30 µM for 48h. After harvesting by trypsinization and washing with PBS, the cells were stained with 5 µL Annexin FITC, incubated for 15 min in the dark and then immediately analyzed using a FACS flow cytometer (BD FACSAria™ II - BD Biosciences) using BD FACSDiva™ Software (BD Biosciences, USA).

### Statistical analysis

The acquired data were analyzed using SPSS 9.0 and the results were given as mean ± SD of three replicates. The mean comparison between the various assessed groups was performed using one-way analysis of variance (ANOVA). Statistical significance was defined when *p* < 0.05.

## Results and discussion

### Lipid production and FA composition

Two oleaginous fungi, i.e. *C. echinulata* and *U. isabellina*, as well as the marine microalgae *N. gaditana*, were selected for this study thanks to their ability to accumulate PUFA-containing lipids.

*C. echinulata* has been recognized as a great GLA producer cultivated in sugar-based media with high C/N ratio [31] while, *U. isabellina* is known for its capability to accumulate lipids in high quantities [32–34]. In this study, *C. echinulata* produced significant quantities of biomass and cellular lipids (i.e. 12.9 g/L of dry biomass containing 30.0% w/w lipids) while *U. isabellina* accumulated 74% of lipids in its dry biomass, both cultivated in a mineral medium with glucose as the sole source of carbon and energy (Table 1). GLA was found in considerable concentration in *C. echinulata* lipids, representing 12.8% of total FAs (Table 2). However, the major FA in these lipids was oleic (C18:1) (with a percentage of 44%), followed by palmitic (C16:0) and linoleic (C18:2) acids. Concerning the FA profile of *U. isabellina*, C18:1 was the dominant FA, found up to 54.4% in total lipids, while C16:0 and C18:2 were also found at significant percentages. GLA percentages were low (i.e. 2.6%) in the lipids of *U. isabellina*. Chatzifragkou *et al*. [35] reported slightly higher GLA percentages in the lipids of both strains compared to the current study. It seems that several factors, such as the carbon source, affect lipid FA composition. For instance, Fakas *et al*. [36], studying the effect of different carbon sources on growth and lipid accumulation rates of *C. echinulata* and *U. isabellina*, reported that xylose, in contrast to glucose, induced lipid accumulation and GLA biosynthesis.

**Table 1.** Biomass yield (x, g or mg/L) and lipid content (L/x, %) of the microorganisms used in this study as source of lipids. The cultures were performed in triplicate

| Microorganism | x | L/x (%) |
| --- | --- | --- |
| *Cunninghamella echinulata* | 12.9 ± 0.9 g/L | 30.0 ± 1.5 |
| *Umbelopsis isabellina* | 13.2 ± 1.2 g/L | 74.0 ± 0.8 |
| *Nannochloropsis gaditana* | 313.9 ± 0.4 mg/L | 22.7 ± 0.1 |

**Table 2.** Fatty acid composition of the methyl ester mixtures used as acyl-donors in the amide synthesis. Analyses were performed in three independent samples

| Source of lipid | Fatty acid composition (% w/w) |  |  |  |  |  |  |  |  |  | Others |
| --- | --- | --- | --- | --- | --- | --- | --- | --- | --- | --- | --- |
|  | C16:0 | C16:1<br>n-7 | C18:0 | C18:1<br>n-9 | C18:2<br>n-6 | C18:3<br>n-6 | C18:3<br>n-3 | C18:4<br>n-3 | C20:1<br>n-9 | C20:5<br>n-3 |  |
| *Cunninghamella* | 15.9 | 2.0 | 7.9 | 44.0 | 13.0 | 12.8 | nd | nd | nd | nd | 4.4 |
| *echinulata* | ± 0.7 | ± 0.5 | ± 0.6 | ± 1.4 | ± 1.2 | ± 1.0 |  |  |  |  | ± 1.2 |
| *Umbelopsis* | 22.1 | 3.6 | 2.8 | 54.4 | 11.7 | 2.6 | nd | nd | nd | nd | 2.8 |
| *isabellina* | ± 1.1 | ± 0.4 | ± 0.4 | ± 4.1 | ± 0.9 | ± 0.3 |  |  |  |  | ± 0.7 |
| *Nannochloropsis* | 18.0 | 20.4 | 0.7 | 13.7 | 5.7 | 0.8 | 0.8 | nd | 7.2 | 25.0 | 7.7 |
| *gaditana* | ± 0.7 | ± 0.5 | ± 0.0 | ± 0.4 | ± 0.1 | ± 0.2 | ± 0.1 |  | ± 0.4 | ± 0.3 | ± 2.7 |
| Olive oil | 12.2 | 2.4 | 2.7 | 74.1 | 7.0 | nd | 0.4 | nd | nd | nd | 1.2 |
|  | ± 1.2 | ± 0.2 | ± 0.3 | ± 1.1 | ± 0.2 |  | ± 0.0 |  |  |  | ± 0.3 |
| EPA concentrate | 0.5 | 0.5 | 3.3 | 10.0 | 1.0 | 0.8 | 2.3 | 2.4 | 4.4 | 72.3 | 4.0 |
|  | ± 0.0 | ± 0.0 | ± 0.2 | ± 1.8 | ± 0.1 | ± 0.0 | ± 0.2 | ± 0.1 | ± 0.8 | ± 1.4 | ± 0.5 |

In the last few decades, the interest in microalgae as PUFA producers is constantly increasing. Biomass production and lipid accumulation of *N. gaditana* cultivated in ASW under constant illumination, were satisfying (i.e. 313.9 mg/L and 22.7%, respectively) and in accordance with data reported by Dourou et al. [22]. The predominant FA was EPA, found at a percentage of 25% in the total FAs, followed by the monounsaturated palmitoleic (C16:1) and C18:1 acids (Table 2). Dourou et al. [22] reported that the same strain cultivated on different bioreactor configurations, was able to synthesize myristic acid (C14:1) too, which can be utilised in a wide variety of applications in the cosmetics industry. The unsaturated FA content, though, could potentially be increased by optimizing the growth conditions and/or by using genetically modified strains.

### Optimization of the FAA synthesis

Initially, the conditions of the amidation reaction (Fig. 1) were optimized by taking the Olive oil FAMEs as a model substrate. The reaction, the progress of which was monitored by TLC, has been done for 24 h at 40 °C with shaking at 90 rpm utilizing acetone as a solvent in the presence of immobilized lipase as a catalyst. The % conversion, which was taken as a criterion for determining the optimum conditions, was quantified via *in situ* NMR monitoring (Fig. 2). Besides, FT-IR analysis (Fig. 3) gave a reliable evidence for amide formation due to the appearance of a band at 1638 cm^−1^, corresponding to the carbonyl of amide, in parallel with the disappearance of the band at 1743 cm^−1^, due to the consumption of the carbonyl group of FAMEs. In addition, amide formation is confirmed by the appearance of a broadband at 3403 cm^−1^ due to NH, which in line with the amide structure 4 and rule out the formation of structure 3 due to the absence of the amino group NH_2_ band (Fig. 1).

**Fig. 1.**
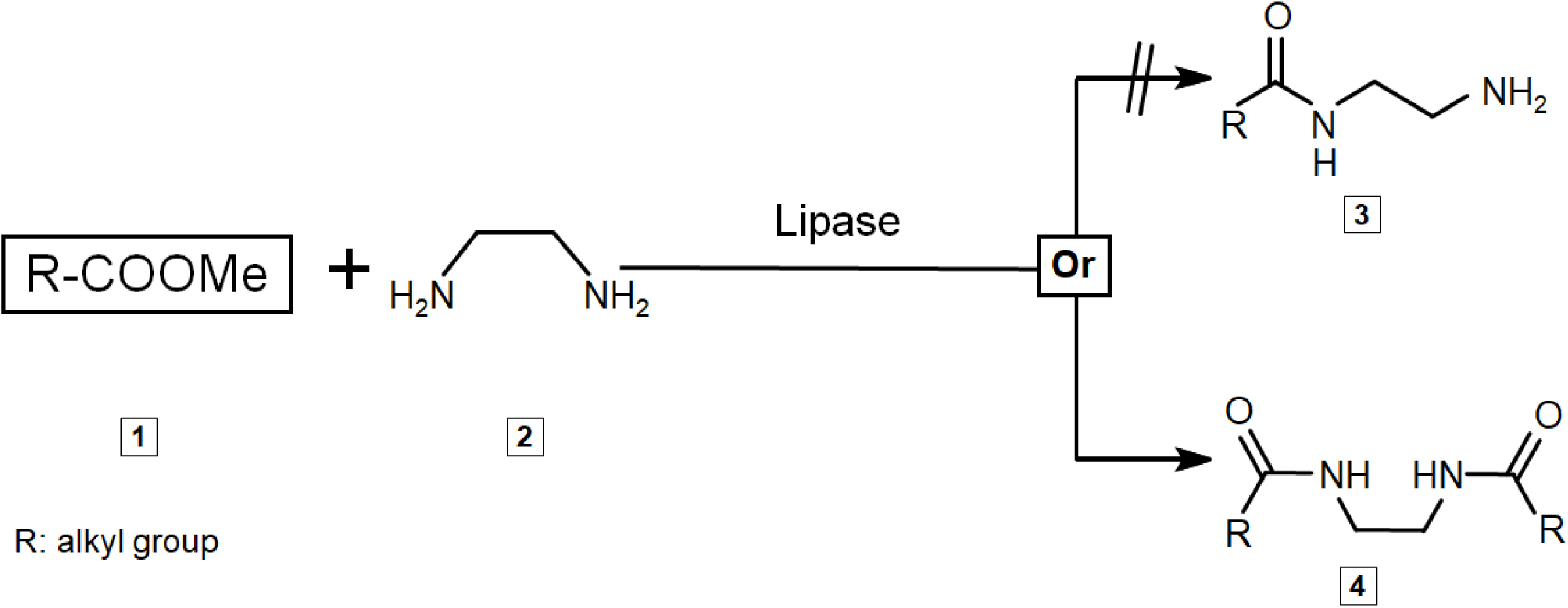
Amidation reaction of ethylene diamine and FAMEs catalyzed by lipase

**Fig. 2.**
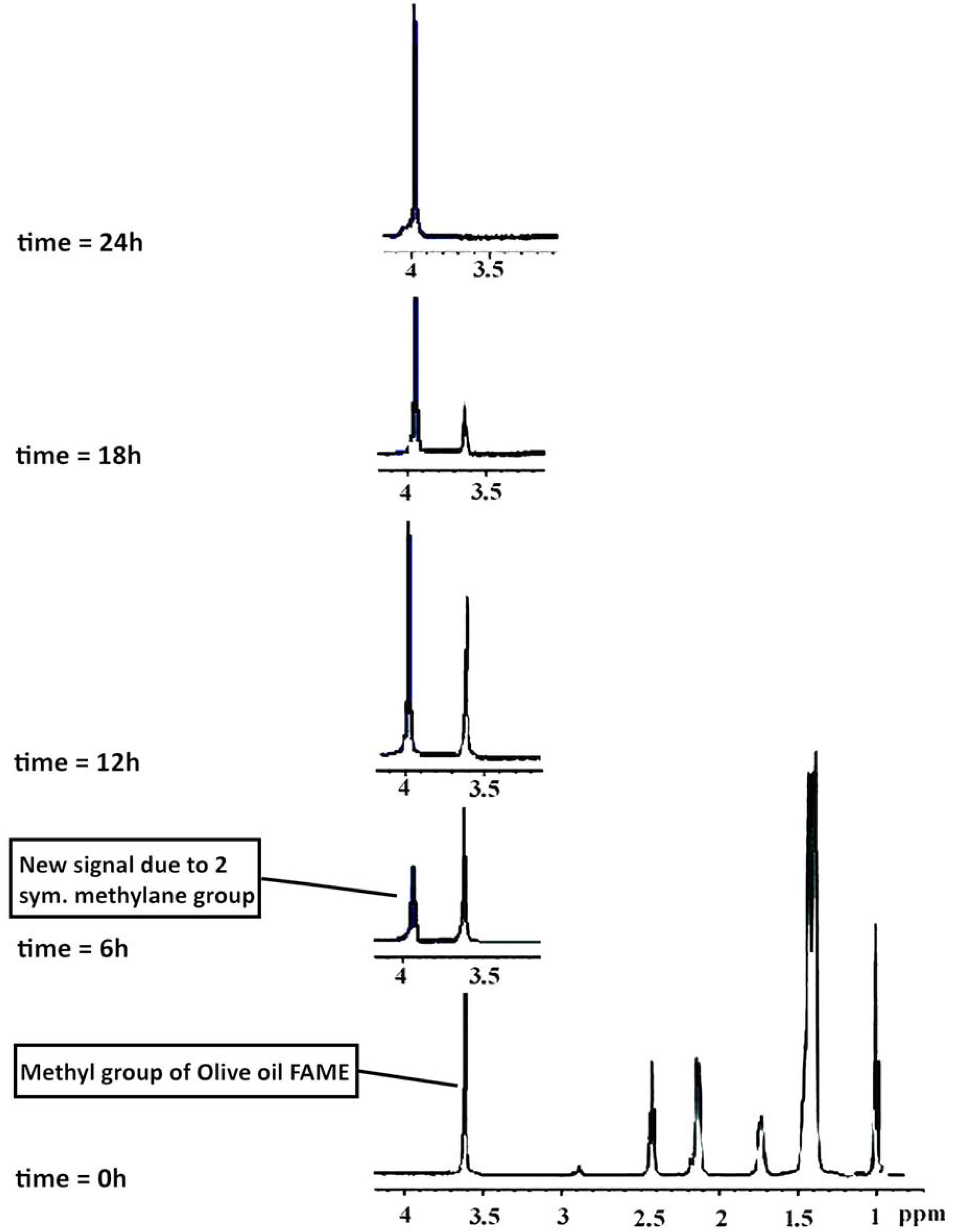
*In situ* NMR monitoring for % conversion of Olive oil FAMEs to amide

**Fig. 3.**
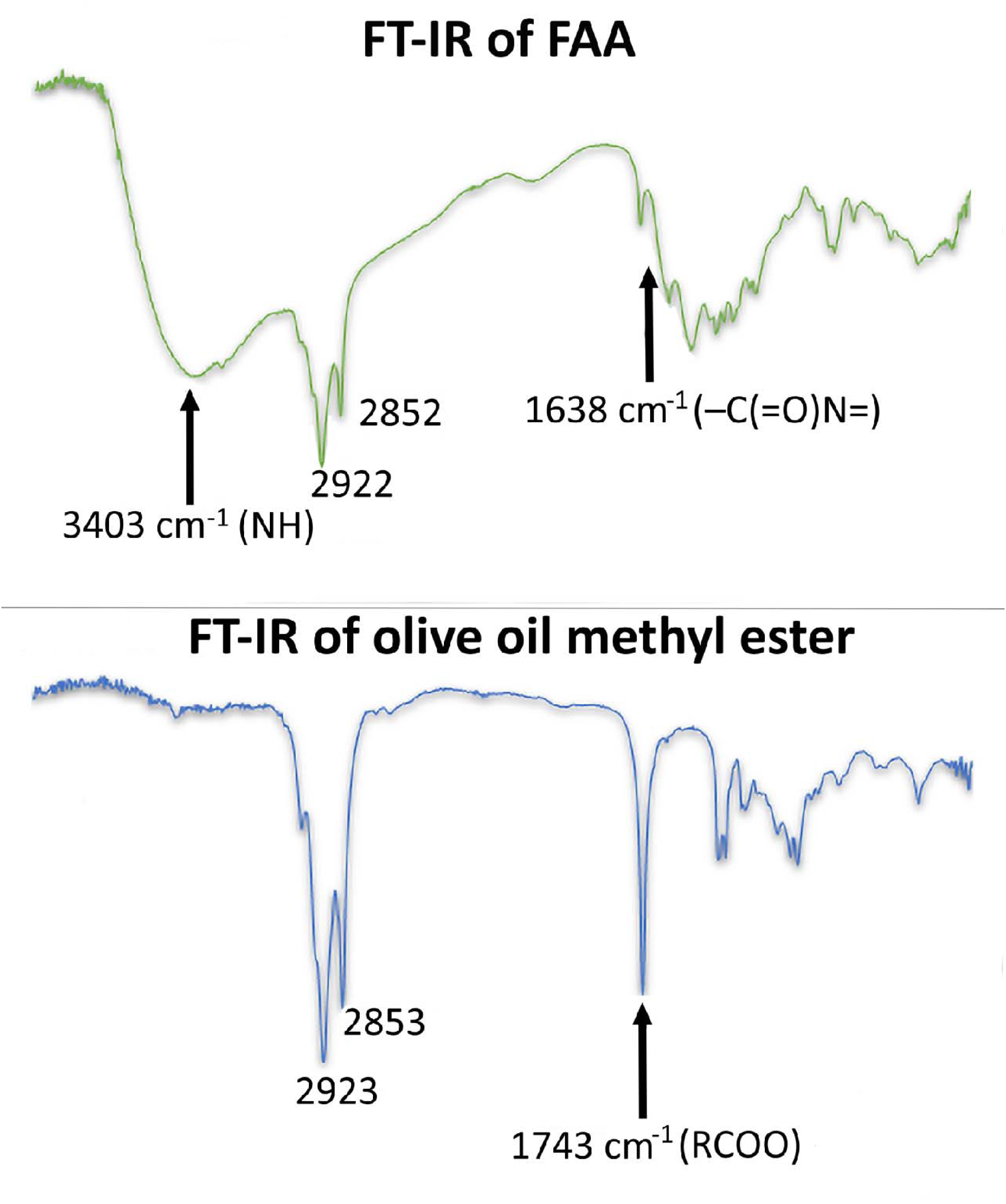
FT-IR analysis of Olive oil FAMEs and its amide

Two immobilized lipases, namely Novozym 435 and lipase from *C. rugosa* (Lipase CR) were used as catalysts for the reaction of Olive oil FAMEs with ethylene diamine (Table 3). The conversion yield was only 12% in the absence of a catalyst (entry 9, Table 3), confirming the importance of lipases in FAA synthesis. Wang et al. [2, 3] proved that even in the absence of a catalyst amidation can be performed but at high temperature and long reaction times, conditions that create undesired product quality. Additionally, it was demonstrated that, under the conditions of the present experimental work, the immobilized lipase Novozym 435 showed lower conversion yield (entries 1-4, Table 3) than the Lipase CR (entries 5-8, Table 3).

**Table 3.** Synthesis of Olive oil-FAAs utilizing different immobilized lipases and different molar ratio of Olive oil FAMEs: ethylene diamine

| Entry | Immobilized enzyme | FAME: Amine Ratio | Conversion (%) |
| --- | --- | --- | --- |
| 1 | Novozym 435 | 1:2 | 31.15 ± 1.25 |
| 2 | Novozym 435 | 1:3 | 57.05 ± 2.95 |
| 3 | Novozym 435 | 1:4 | 76.30 ± 3.30 |
| 4 | Novozym 435 | 1:5 | 95.00 ± 4.15 |
| 5 | Lipase CR | 1:2 | 39.02 ± 2.95 |
| 6 | Lipase CR | 1:3 | 71.11 ± 4.55 |
| 7 | Lipase CR | 1:4 | 89.50 ± 3.45 |
| 8 | Lipase CR | 1:5 | 100.00 ± 4.83 |
| 9 | No catalyst | 1:5 | 12.57 ± 0.43 |

Immobilized enzymes have the advantage over free enzymes to be easily recycled, providing sustainability to the process, and for this reason several researchers proposed the employment of immobilized lipases as catalysts for FAA synthesis [37–39]. In the current work the reusability of the Lipase CR was checked for several reaction cycles for the synthesis of amide of Olive oil FAMEs under the optimized reaction conditions. Specifically, the enzyme was removed after the completion of the reaction by filtration, washed with ethanol solvent in a Soxhlet extraction apparatus and the recovered enzyme was reused for three times under the same reaction conditions. It was found that the regenerated enzyme performed the reaction efficiently without loss of its catalytic activity. Similarly, according to Khare et al. [8], Novozym 435 could be repeatedly used without any decrease of its catalytic activity, while Sharma et al. [40] have demonstrated six repeated cycles of reusability of Chirazyme L-2 used to synthesize secondary amide surfactant from N-methylethanol amine.

Previously, Wang et al. [2] showed that the molar ratio of vinyl stearate to ethanolamine has affected the synthesis and purity of N-stearoyl ethanolamine produced. Thus, in the present research different molar ratios of Olive oil FAMEs: ethylene diamine were tested (Table 3, Fig. 4). The amine rather than the FAME concentration in the reaction medium affected the conversion yield of Olive oil FAMEs to amide. The maximum conversion yield (i.e. 100 %) was attained with Lipase CR and a ratio of Olive oil FAMEs: ethylene diamine 1:5 (Table 3, entry 8). These results are almost similar with those reported by Liu et al. [41] who, studying the effect of the FA: diethanolamine ratio on the lipase-catalyzed amidation, showed that the maximum amide yield was achieved at a low ratio 1:4.

**Fig. 4.**
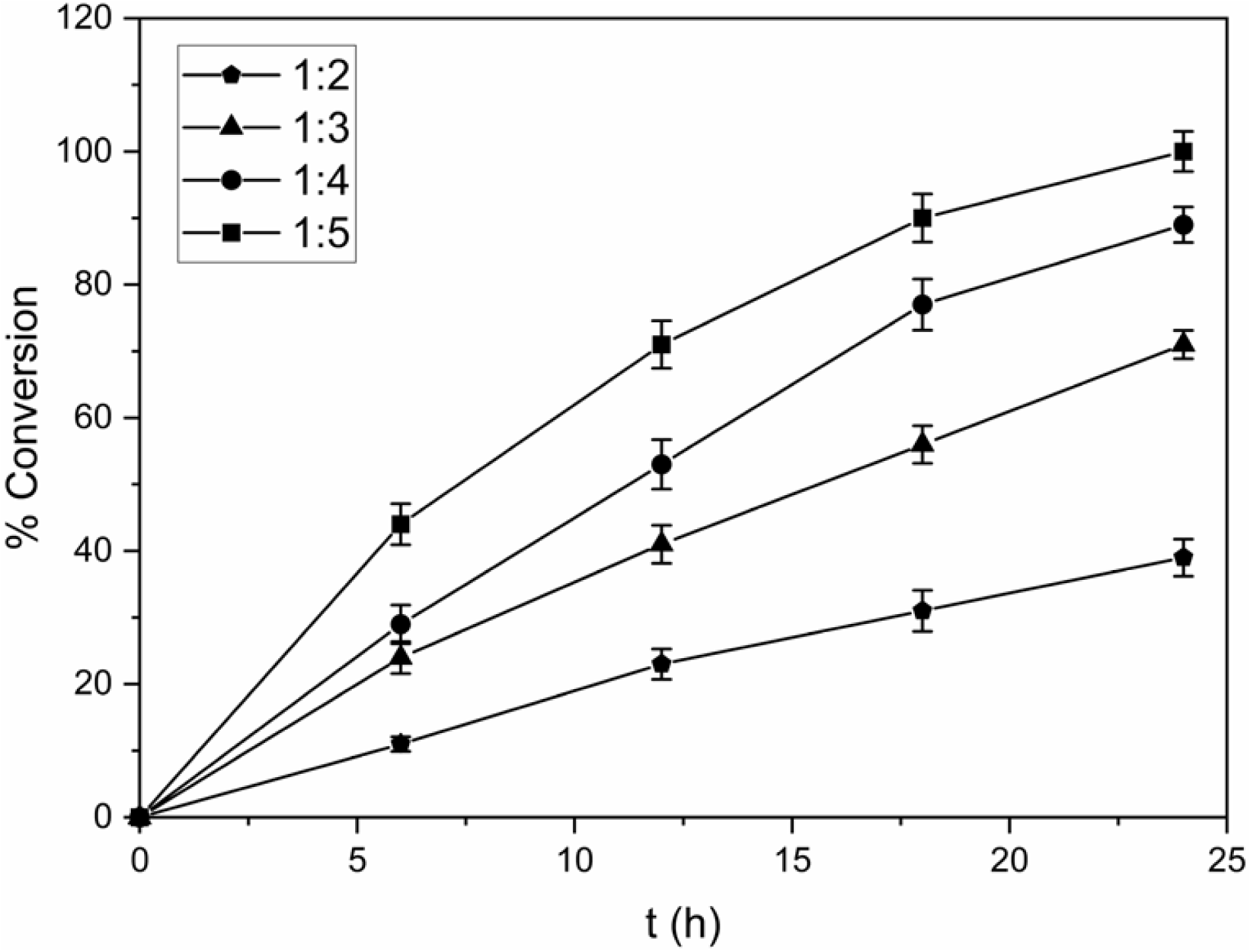
% conversion of Olive oil FAMEs to amide in different ratios of Olive oil FAMEs: ethylene diamine

The reaction conditions were further optimized using the immobilized Lipase CR as a catalyst in different quantities and the reaction progress was monitored by TLC. It was found that a 100% conversion yield was obtained in the shortest reaction time when 0.1 g of the Lipase CR was employed (Table 4, entry 2), while a higher enzyme quantity was not necessary. Wang et al. [3] reported that the yield of the amidation reaction increased when lipase concentration increased from 10 to 20% of the total reactants.

**Table 4.** Optimization of reaction conditions for Olive oil-FAAs synthesis utilizing Lipase CR

| Entry | Lipase CR<br>(wt, g) | Solvent | Time<br>(h) | Conversion<br>(%) |
| --- | --- | --- | --- | --- |
| 1 | 0.05 | Acetone | 30 | $82.09 \pm 2.15$ |
| 2 | 0.10 | Acetone | 24 | $100.04 \pm 3.48$ |
| 3 | 0.15 | Acetone | 24 | $99.13 \pm 4.22$ |
| 4 | 0.10 | t-butyl alcohol | 30 | $98.95 \pm 4.78$ |
| 5 | 0.10 | Iso-amyl alcohol | 30 | $100.20 \pm 3.45$ |

Due to the nature of the reactants organic solvents were used by many researchers as a suitable medium for the production of FAAs (see for instance [41, 42]). In the current paper when acetone was used as a solvent the duration of the reaction was shorter than when *t-*butyl alcohol and isoamyl alcohol were employed (entries 2, 4, 5, Table 4). Acetone is an environmentally benign and low toxicity solvent, previously used as the best solvent for polyunsaturated FAA synthesis [43].

Following optimization of the amidation reaction FAAs were produced using FAMEs of SCOs from *C. echinulata, U. isabellina* and *N. gaditana* and of EPA concentrate oil (Table 5). The conversion yields of the above reactions were excellent, reaching the values of 90-100% (Table 5). The structures of the obtained FAAs were confirmed on the basis of their FT-IR spectra in which appearance of the broadband due to NH group and of carbonyl group of amides was observed (see for instance Fig. 5, 6 and original spectra in Fig. S1-S5). These results are in agreement with Mudiyanselage et al. [44] who showed that microalgal lipids can be converted into FAAs in a two-step reaction, including transesterification to form FAMEs followed by amidation. The synthesis of amide in this work is efficient and its potential application on a large scale will not interfere with the food supply chain, since SCOs are alternative sources to the traditional sources of PUFAs.

**Table 5.** Synthesis of FAAs by Lipase CR in acetone medium utilizing FAMEs of different origin

| Entry | Source of FAMES | Time<br>(h) | Conversion<br>(%) |
| --- | --- | --- | --- |
| 1 | *Cunninghamella echinulata* | 24 | 90.23 ± 2.51 |
| 2 | *Umbelopsis isabellina* | 18 | 95.36 ± 5.75 |
| 3 | *Nannochloropsis gaditana* | 18 | 90.95 ± 1.50 |
| 4 | EPA concentrate | 18 | 100.05 ± 3.00 |

**Fig. 5.**
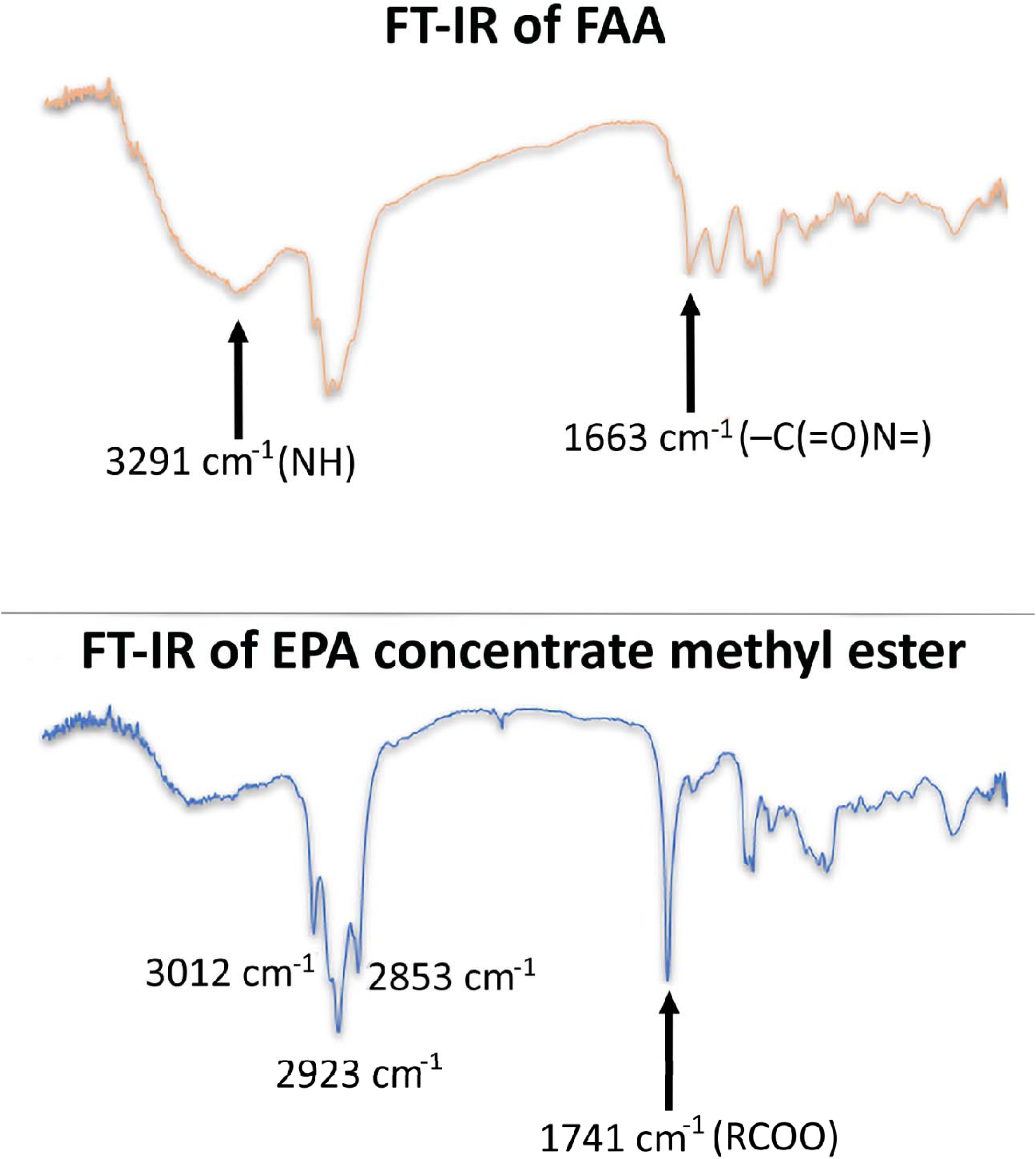
FT-IR analysis of EPA concentrate FAMEs and its amide

**Fig. 6.**
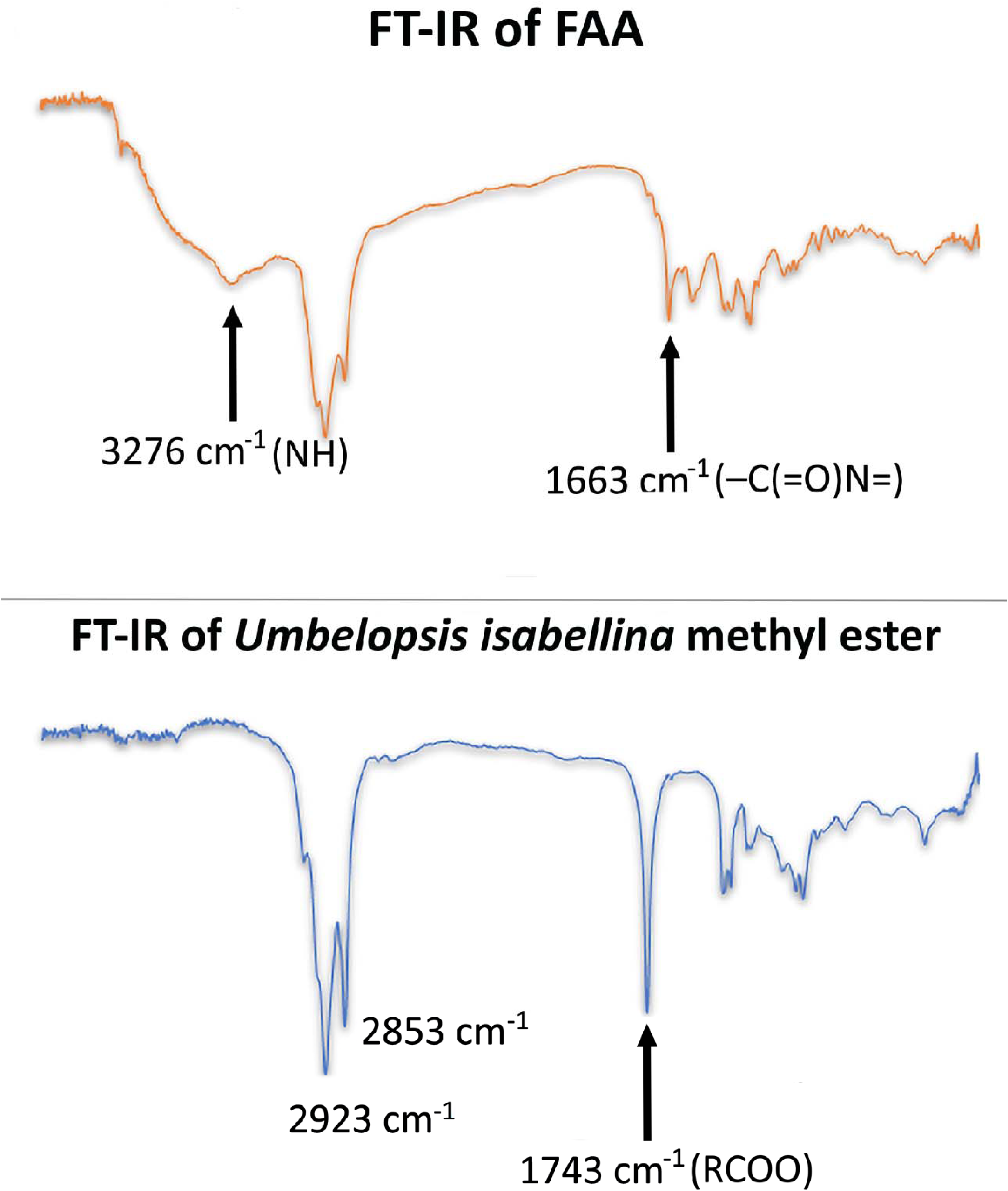
FT-IR analysis of *Umbelopsis isabellina* FAMEs and its amide

### Antimicrobial activity of amide compounds

FAAs derived from FAMEs of *C. echinulata, U. isabellina, N. gaditana* SCOs, Olive oil and EPA concentrate were tested against various human pathogens for their antimicrobial efficacy by the agar well diffusion method, which resulted in the formation of a variable diameter zone of inhibition (Table 6). Except for MRSA *Staphylococcus aureus*, which is inhibited only by *N. gaditana*-FAAs, Olive oil-FAAs and EPA-FAAs, all tested pathogens were significantly inhibited by all FAAs produced in this work. *U. isabellina*-FAA was probably the most efficient preparation against all pathogens, except for *MRSA Staphylococcus aureus*. On the contrary, *C. echinulata*-FAAs seemed to be less efficient than *U. isabellina*-FAAs against all pathogens (statistically significant at p<0.05). *N. gaditana*-FAAs successfully inhibited all tested organisms, except for *Bacillus subtilis* ATCC 6633 in the culture of which the inhibition zone was only 9.00 ± 0.00 mm. The inhibition demonstrated by *N. gaditana*-FAAs against *Staphylococcus aureus* ATCC 25923 and *Candida albicans* ATCC 10221 was similar to that observed when *U. isabellina*-FAAs were employed. The Olive oil-FAAs showed a significant inhibitory activity against all tested organisms, especially against *Pseudomonas aeruginosa* ATCC 15442 and *Candida albicans* ATCC 10221 presenting an inhibition zone, 17.67 ± 0.57 mm and 18.07 ± 0.11 mm, respectively. Finally, EPA-FAAs showed a high antimicrobial activity against all tested organisms specifically against *Staphylococcus aureus* (both strains) and *Pseudomonas aeruginosa* ATCC 15442 (i.e. inhibition zone around 20 mm).

**Table 6.** The antimicrobial activity of FAAs against pathogenic strains. Data represent the mean of the diameter of the inhibition zones of three replicates ±SD. The concentration of FAAs used to determine the diameter of the inhibition zones was 4 mg/mL

|  | Source of FAAs |  |  |  |  |
| --- | --- | --- | --- | --- | --- |
|  | *Cunninghamella echinulata* | *Umbelopsis isabellina* | *Nannochloropsis gaditana* | Olive oil | EPA concentrate |
|  | Inhibition zone (mm) |  |  |  |  |
| *Escherichia coli*<br>(ATCC 25922) | 9.00 ± 1.00 | 17.00 ± 0.00 | 16.1 ± 1.01 | 15.67 ± 1.15 | 14.0 ± 0.00 |
| *Klebsiella pneumoniae*<br>(ATCC 700603) | 10.33 ± 0.58 | 16.00 ± 0.00 | 14.3 ± 0.15 | 13.57 ± 0.81 | 12.57 ± 0.11 |
| *Salmonella typhimurium*<br>(ATCC 14028) | 9.33 ± 0.58 | 19.00 ± 0.0 | 17.33 ± 0.57 | 14.67 ± 1.15 | 16.00 ± 0.00 |
| *Pseudomonas aeruginosa*<br>(ATCC 15442) | 12.00 ± 1.00 | 20.00 ± 0.5 | 18.00 ± 0.00 | 17.67 ± 0.57 | 20.00 ± 1.00 |
| *Bacillus subtilis*<br>(ATCC 6633) | 14.17 ± 0.29 | 17.00 ± 0.17 | 9.00 ± 0.00 | 12.17 ± 1.28 | 17.00 ± 1.00 |
| MRSA *Staphylococcus aureus*<br>(ATCC 4330) | 0.00 ± 0.00 | 0.00 ± 0.00 | 15.50 ± 0.50 | 13.67 ± 1.15 | 19.00 ± 1.00 |
| *Staphylococcus aureus*<br>(ATCC 25923) | 9.17 ± 0.29 | 22.00 ± 0.5 | 21.00 ± 0.86 | 15.00 ± 0.00 | 20.00 ± 2.00 |
| *Candida albicans*<br>(ATCC 10231) | 16.00 ± 2.00 | 21.00 ± 1.8 | 19.00 ± 0.5 | 18.07 ± 0.11 | 18.07 ± 0.11 |

The results of MIC and MBC determined for selected pathogens (Table 7) were in line with those obtained by the agar diffusion method. In detail, all tested pathogenic strains are sensitive to the *U. isabellina*-FAAs, while *C. echinulata*-FAAs are less effective. For *N. gaditana*-FAAs, the highest MIC observed was 200 µg/mL and this was against *Bacillus subtilis* ATCC 6633, while the other pathogens tested were much more sensitive. All strains are sensitive to FAAs derived from Olive oil, especially *Bacillus subtilis* ATCC 6633 and *Pseudomonas aeruginosa* ATCC 15442 in which MIC was only 25 µg/mL. Besides, the FAAs derived from EPA concentrate significantly affected the growth of all the tested bacteria particularly of *Bacillus subtilis* ATCC 6633 and *Klebsiella pneumoniae* ATCC 700603. With few exceptions (case of EPA-FAAs) the MBC was estimated to be 100-200 µg/mL.

**Table 7.**
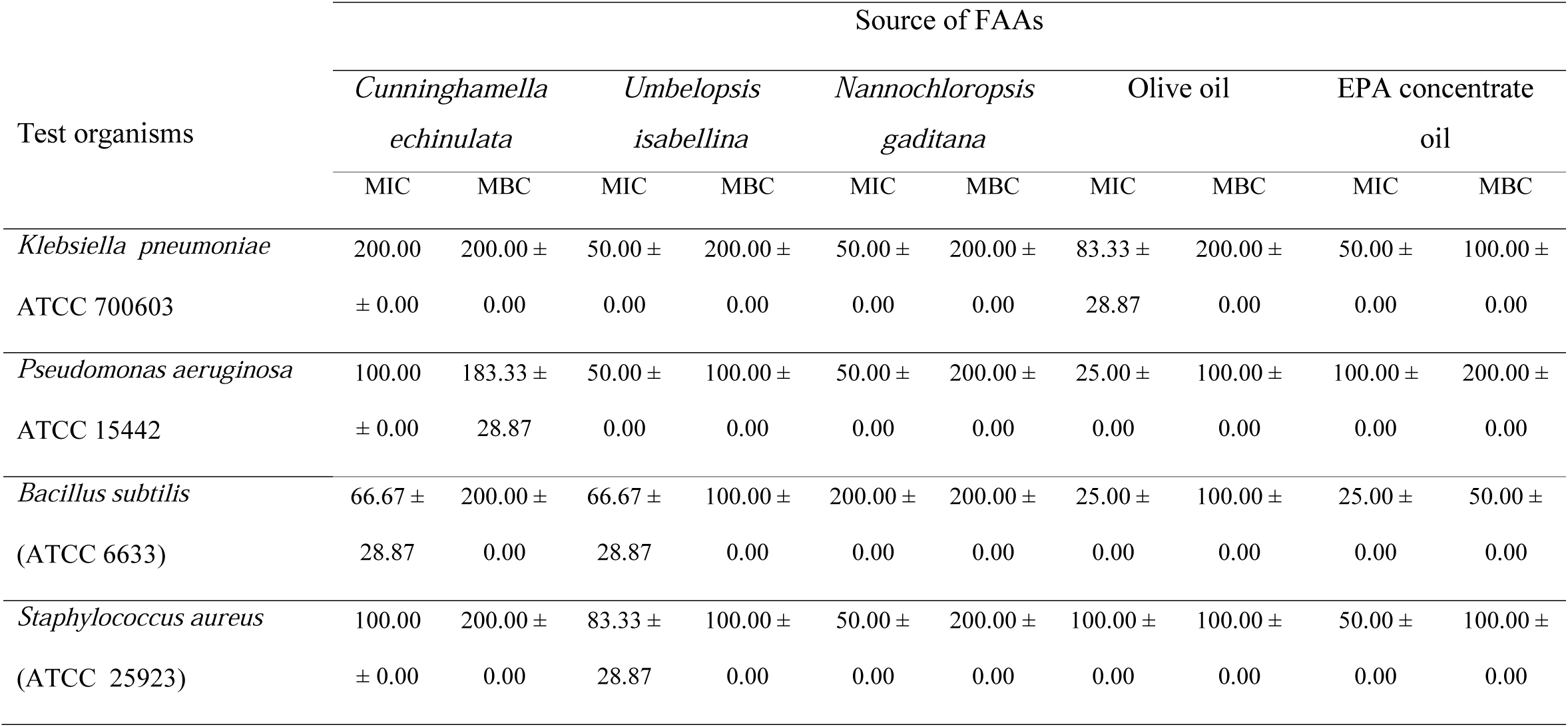
Determination of minimum inhibitory concentration (MIC, µg/mL) and minimum bactericidal concentration (MBC, µg/mL) of FAAs against pathogenic strains

The results reported in this paper are in agreement with previous reports in which various FAAs have been used as potential antimicrobial agents [7]. Khare et al. [8] showed that FAAs possess a strong antimicrobial activity towards Gram-positive (such as *Bacillus subtilis* and *Staphylococcus aureus*) and Gram-negative (such as *Proteus vulgaris* and *Klebsiella pneumoniae*) bacteria. Concerning the mechanism of action of FAAs Novak et al. [45] observed that FAAs containing an epoxy group exhibit a broad spectrum of antimicrobial activity, which is further enhanced by unsaturation. Later, Stevens and Hofmeyr [46] indicated a disturbance in the FA constituents of the cell plasma membrane which interferes with the proper membrane functions leading to the loss of the integrity of the plasma lemma. This suggestion was further strengthened by Shao *et al*. [47] who worked on the mechanism of action of oleamide observed that this compound caused alterations in the FA composition of the cell membranes. In this paper FAAs prepared using FAMEs rich in oleic acid (i.e. Olive oil-FAAs and *U. isabellina*-FAAs) were more effective against pathogens than those prepared from FAMEs contained oleic acid in lower concentration (i.e. *C. echinulata*-FAAs). Unexpectedly, the presence of GLA in the lipids of *C. echinulata* did not improve the antimicrobial activity of *C. echinulata*-FAAs compared to *U. isabellina*-FAAs. *N. gaditana*-FAAs and EPA-FAAs, although containing oleic acid in very low concentrations, are both effective against all tested bacteria, and therefore their activity may be attributed to their high EPA content.

### Insecticidal activity assay of amide compounds

The yellow fever mosquito (*Aedes aegypti*) spreads dangerous human arboviruses that include dengue, Zika, and chikungunya. Therefore, control of yellow fever mosquitoes is a critical public health priority [48]. Chemical insecticides are a leading method of control, but they are expensive, contribute to the development of insecticidal resistance, pose risks to the environment and cause safety concerns to humans and non-target animal species.

The susceptibility of *Aedes aegypti* larvae to FAAs under laboratory conditions was tested using dipping methods. The larvicidal activity of a product is usually improved by increasing its concentration and exposure time [49]. Many FA-derived products manifest toxicity to different mosquito species larvae [50, 51], and have been proposed as alternatives to conventional mosquito larvicides. Komalamisra et al. [52] considered larvicidal compounds exerting LC50□<□50 mg/L active, 50 mg/L<□LC50□<□100 mg/L moderately active, 100 mg/L <□LC50□<□750 mg/L effective, and LC50□>□750 mg/L inactive. Kiran et al. [53] considered compounds with LC50□<□100 mg/L as exhibiting a significant larvicidal effect. In the current study, *C. echinulata*-FAAs showed a strong insecticidal activity against *A. aegypti* larvae with LC50 0.3 mg/L, which could be probably attributed to the presence of GLA in significant concentrations, while Olive oil-FAAs, EPA-FAAs and *N. gaditana*-FAAs exhibited active insecticidal effect, demonstrating LC50 18.3, 20.5 and 34.3 mg/L, respectively. Contrary, *U. isabellina*-FAAs were less active, presenting a LC50 equal to 132.1 mg/L (Table 8, Fig. 7). Many bioactive substances, such as plant essential oils [54, 55], FAs [56] and cyanobacterial extracts [57, 58], have been tested against *A. aegypti* larvae. According to our knowledge, this is the first report demonstrating a larvicidal activity of FAAs, in contrast with numerous works dealing with plant-based derivatives [59].

**Table 8.** Susceptibility of *Aedes aegypti* larvae to FAAs under laboratory conditions by using dipping methods. Data represent the mean of six replicates

| Source of FAAs | LC50<br>(mg/L) | Lower<br>limit | Upper<br>limit | RR |
| --- | --- | --- | --- | --- |
| *Cunninghamella echinulata* | 0.294 | 0.258 | 0.332 | 1.00 |
| *Umbelopsis isabellina* | 38.837 | 34.29 | 43.588 | 132.10 |
| *Nannochloropsis gaditana* | 10.073 | 5.481 | 23.647 | 34.26 |
| Olive oil | 5.391 | 4.228 | 6.685 | 18.34 |
| EPA concentrate | 6.039 | 2.592 | 10.733 | 20.54 |

**Fig. 7.**
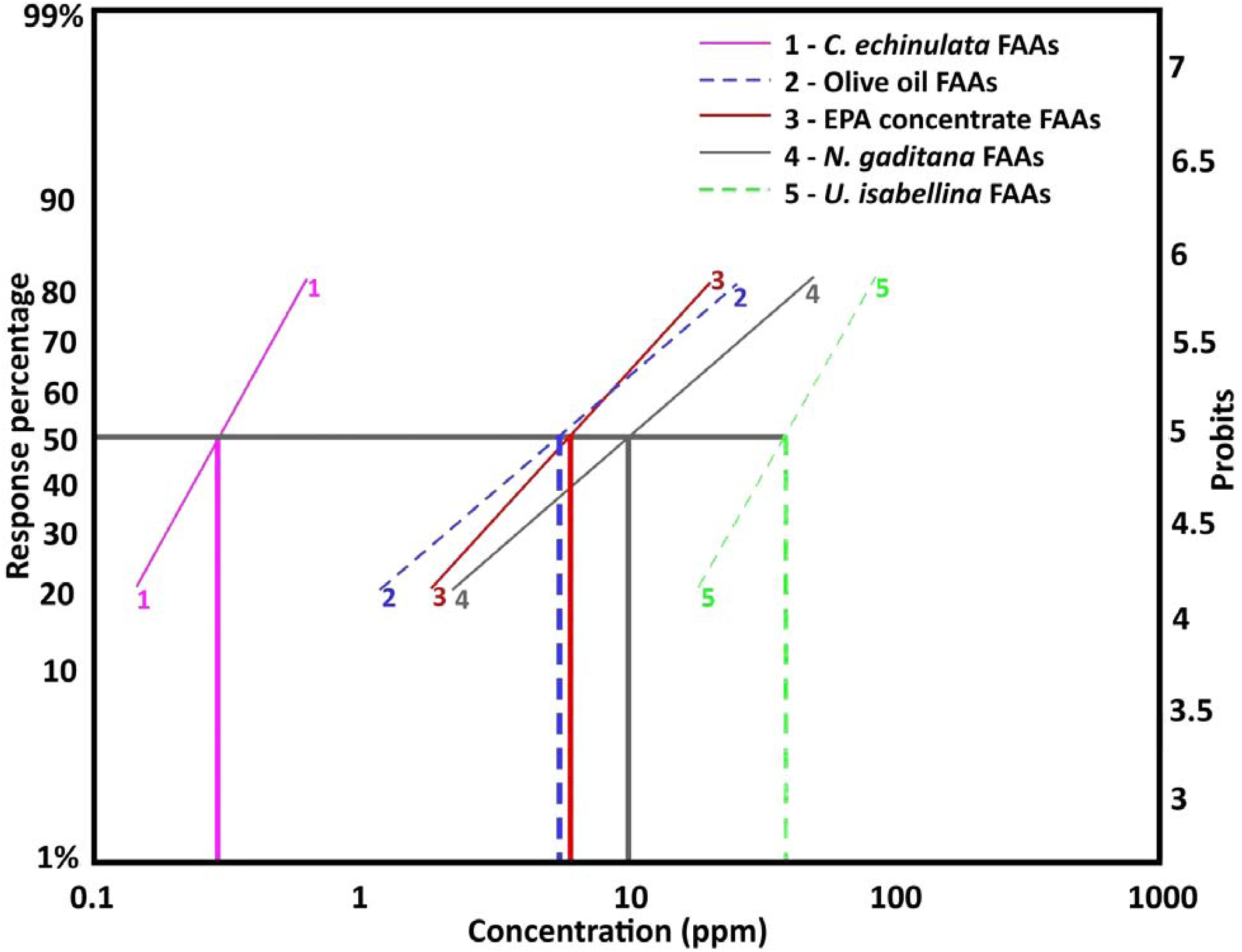
The larval mortality effect of FAAs of *C echinulata, U isabellina, N gaditana*, Olive oil and EPA concentrate at different concentrations against *Aedes aegypti* after continuous exposure for 48 hours

### Cell apoptosis of ovarian cancer cell line induced by FAAs

The results show that all FAAs produced in this work can induce apoptosis of the SKOV-3 ovarian cancer cell line. Higher percentage of apoptosis was observed in the cells treated with *N. gaditana*-FAAs followed by EPA-FAAs, Olive oil-FAAs and *C. echinulata*-FAAs (i.e. 61.7, 54.7, 52.7 and 50.4, respectively). The results are presented in Fig. 8.

**Fig. 8.**
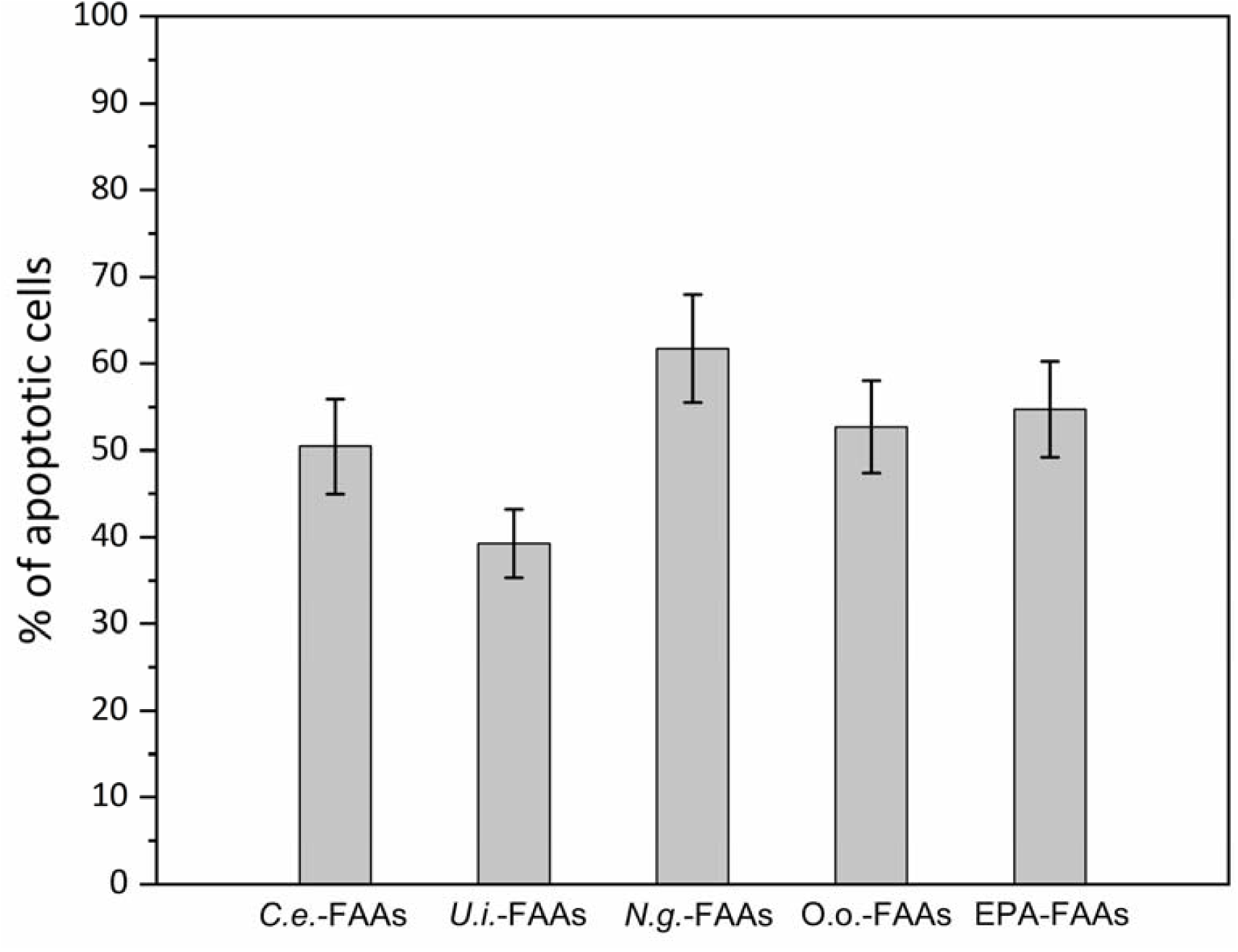
Effect of compounds on SKOV-3 cell apoptosis. Flow cytometry analysis of apoptosis in SKOV-3 cells either untreated or treated with 10 µg/mL of every compounds for 48h. After the treatment period, the cells were stained with Annexin FITC and subsequently analyzed by flow cytometry **Abbreviations:** *C. echinulata* – FAAs: *C*.*e*. – FAAs; *U. isabellina* – FAAs: *U*.*i*. – FAAs; *N. gaditana* – FAAs: *N*.*g*. – FAAs; Olive oil – FAAs; O.o. – FAAs; EPA concentrate – FAAs: EPA – FAAs

Santos et al. [60] suggested that the antiproliferative activity is influenced by the structural variation in the FAAs. Several studies have shown that intake of EPA, the main FA of *Nannochloropsis* sp. may play a role in the prevention of the development of different type of cancer [61]. Particularly, EPA and DHA have been investigated as potential dietary-based agents for breast cancer prevention [62], and they have been shown to exhibit multiple anticancer mechanisms of action, including the alteration of cell signaling [63], inhibition of cell proliferation [64], inflammation [65], metastasis [64, 65], as well as induction of apoptosis [64, 66]. Studies with synthetic FAAs showed antiproliferative activity against several tumor cells [67] and therefore these amide mediators may provide promising new agents, active against inflammatory and cancer diseases [68]. It seems that variation in the FA moieties on groups attached to the nitrogen atom may be responsible for differences in antiproliferative profiles [69].

## Conclusions

FAMEs derived from SCOs of different origin and FA composition can be used in FAA (diamide) enzymatic synthesis catalyzed by immobilized lipases, such as the *Candida rugosa* lipase. The reaction of FAA synthesis can be completed under environmentally friendly conditions in 24 h, while both the solvent (acetone) and the enzyme can be recycled. The biological activities (antimicrobial, insecticidal activity, anti-cancer) of the synthesized FAAs are related, partially at least, to their FA profile. Therefore, we conclude that oleaginous microorganisms, able of synthesizing a wide range of FAs, can be considered in the near future as source FAs suitable for producing FAAs of different biological activities.

## Supporting information

Original spectra of FT-IR analysis on Supplemental Figures S1-S5

## Abbreviations

ANOVA: Analysis of variance
ASW: Artificial sea water
CLSI: Clinical and Laboratory Standards Institute
DHA: Docosahexaenoic acid
EPA: Eicosapentaenoic acid
FA: Fatty acid
FAAs: Fatty acid amides
FAMEs: Fatty acid methyl esters
FT-IR: Fourier-transform infrared
GLA: Gamma linolenic acid
LC50: Median lethal concentration
MBC: Minimum bactericidal concentration
MHA: Mueller Hinton II Agar
MIC: Minimum inhibitory concentration
NMR: Nuclear magnetic resonance
OPSR: Open-pond simulating reactor
PDA: Potato dextrose agar
PUFAs: Polyunsaturated fatty acids
SCOs: Single cell oils
TLC: Thin-layer chromatography

## Author agreement

Hatim A. El-Baz, Ahmed M. Elazzazy, Tamer S. Saleh, Panagiotis Dritsas, Jazem A. Mahyoub, Mohammed N. Baeshen, Hekmat R. Madian, Mohammed Alkhaled and George Aggelis have all agreed to submission.

## Acknowledgments

This work was funded by the *University of Jeddah*, Saudi Arabia, under grant No. (UJ-06-18-ICP). The authors, therefore, acknowledge with thanks the University technical and financial support.

