## Supplementary material for "Enzymatic synthesis of fatty acid amides using microbial lipids as acyl group-donors and their biological activities": Original spectra of FT-IR analysis on Supplemental Figures S1-S5

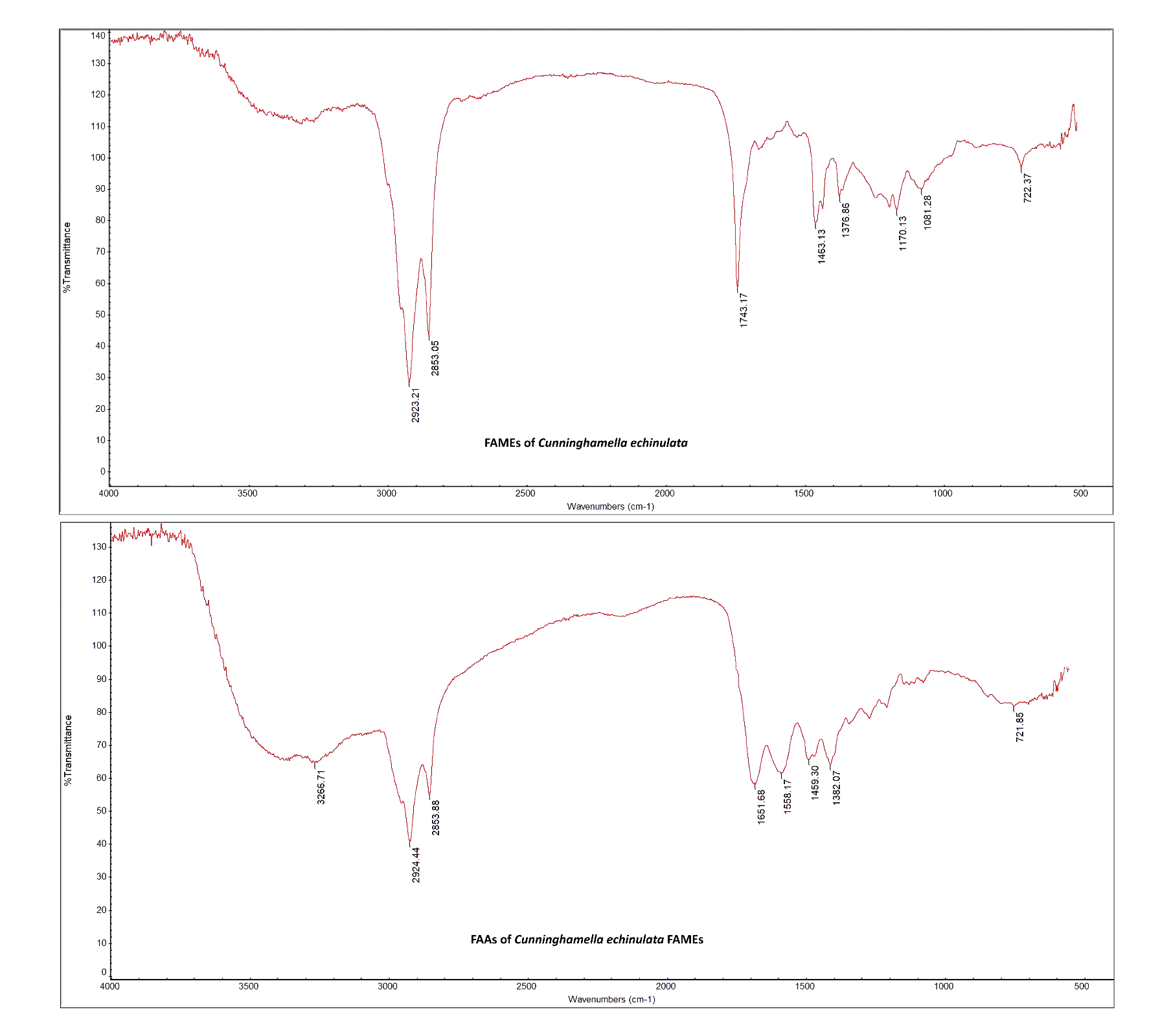
**Fig. S1** FT-IR analysis of *Cunninghamella echinulata* FAMEs and its amide

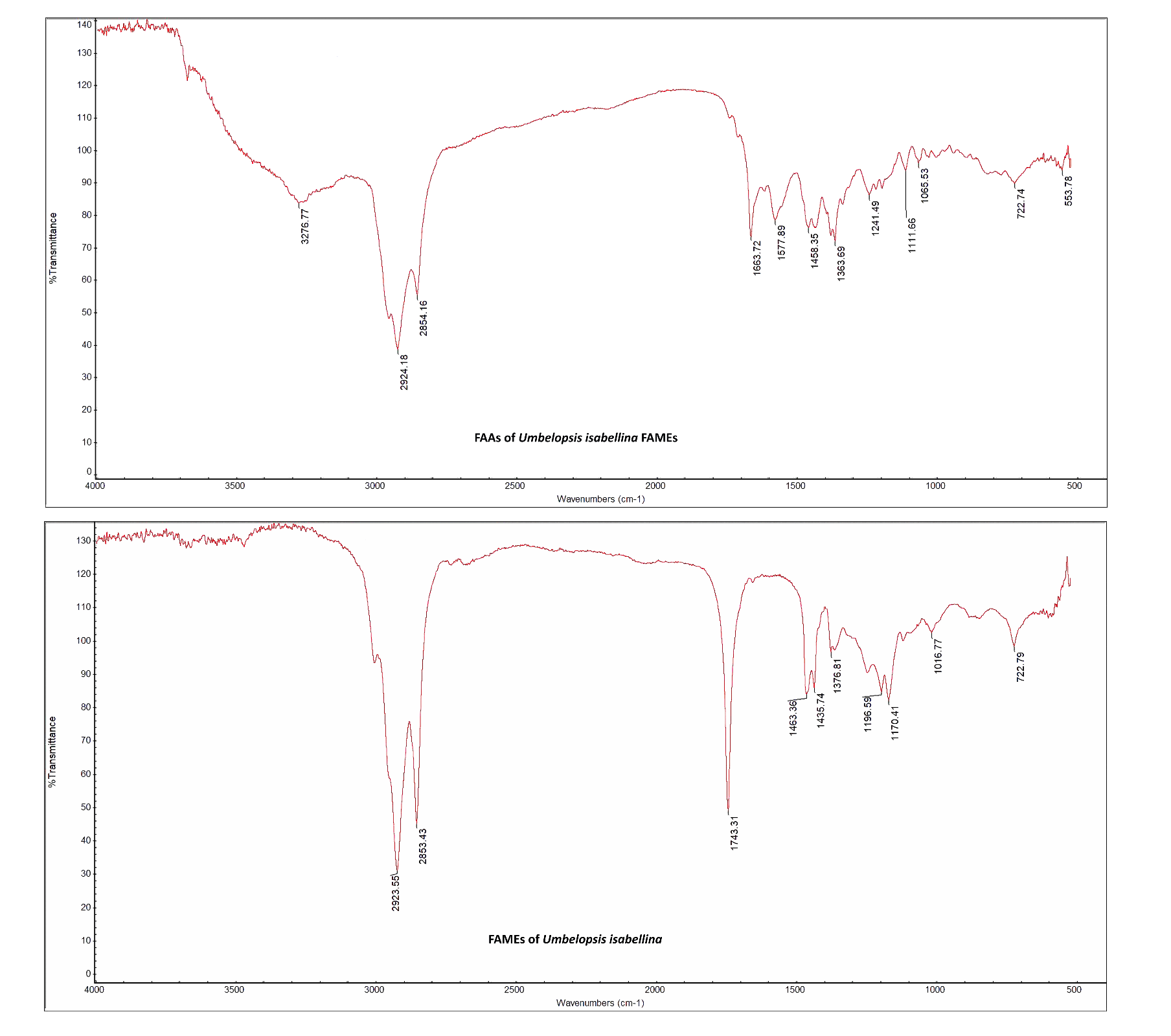
**Fig. S2** FT-IR analysis of *Umbelopsis isabellina* FAMEs and its amide

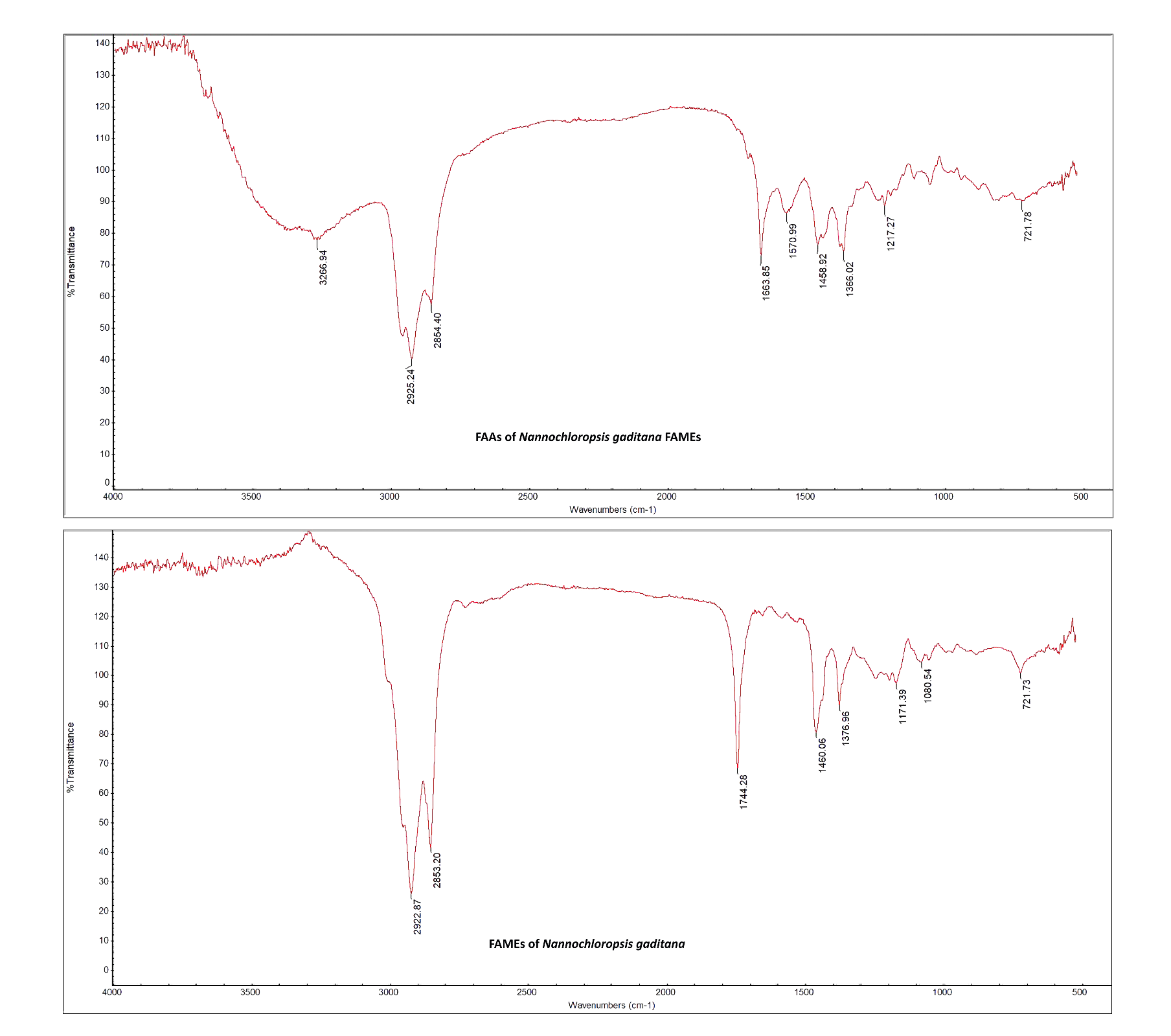
**Fig. S3** FT-IR analysis of *Nannochloropsis gaditana* FAMEs and its amide

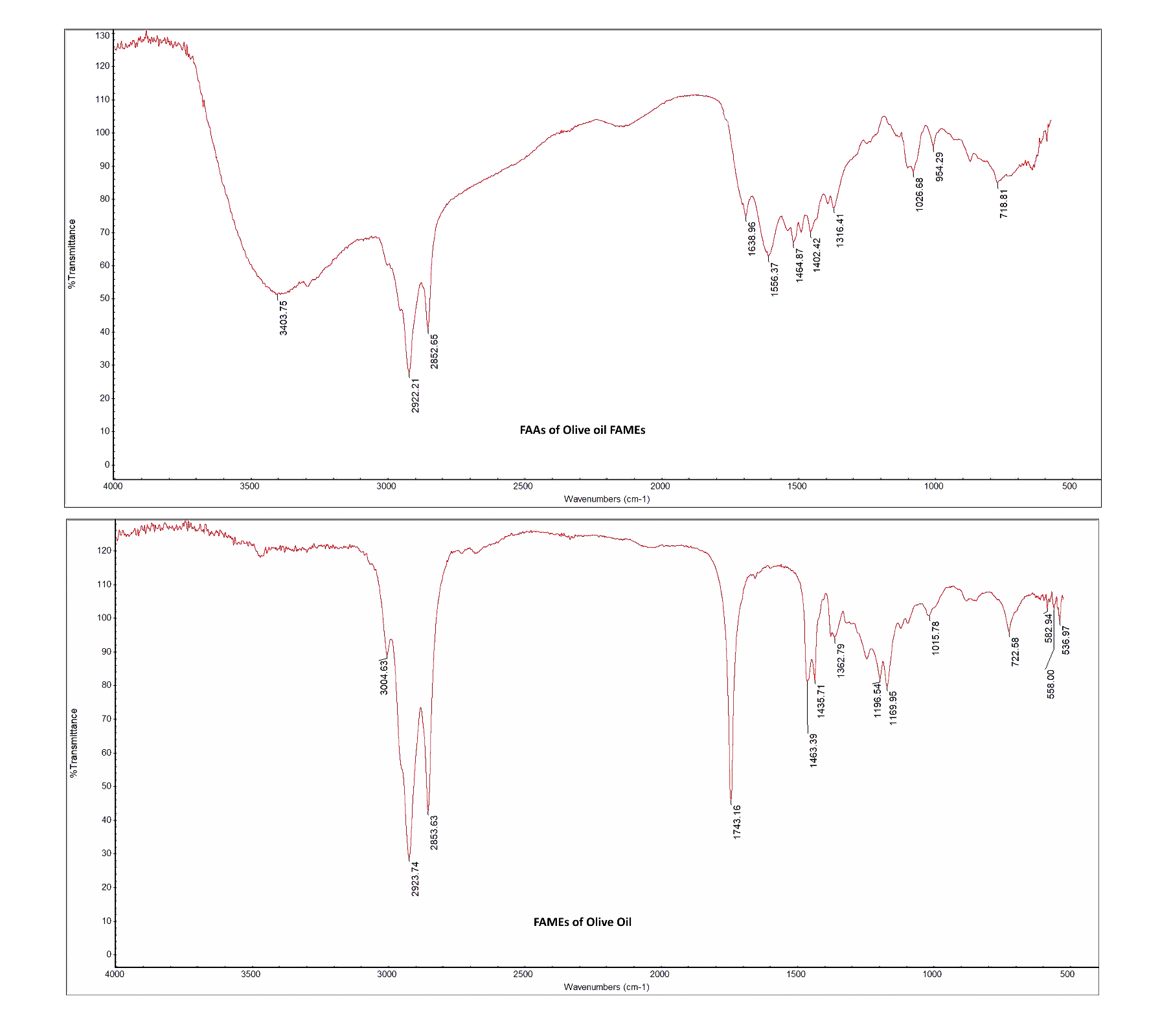
**Fig. S4** FT-IR analysis of Olive oil FAMEs and its amide

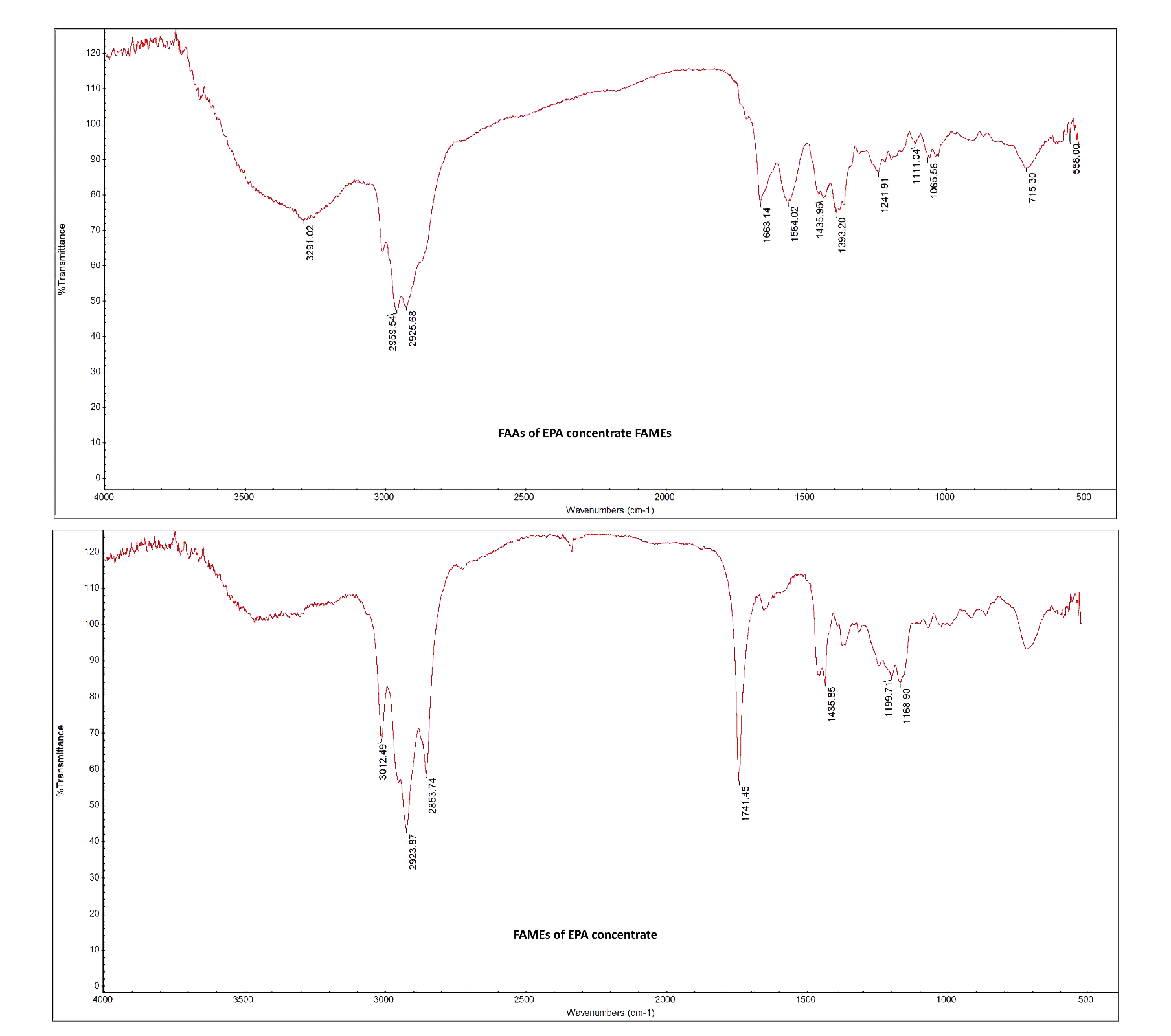
**Fig. S5** FT-IR analysis of EPA concentrate FAMEs and its amide

The original spectra of FT-IR analysis of *Cunninghamella echinulata*, *Umbelopsis isabellina*, *Nannochloropsis gaditana*, Olive oil and EPA concentrate FAMEs and their amides are presented in the Figs. S1-S5.
